# The tumour suppressor Fat1 is dispensable for normal murine hematopoiesis

**DOI:** 10.1101/2023.12.20.572284

**Authors:** Qing Zhang, Meng Ke Li, Xin Yuan Hu, Ying Ying Wang, Pan Pan Zhao, Lin Na Cheng, Rong Hua Yu, Xu Dong Zhang, Song Chen, Zun Min Zhu, Charles E. de Bock, Rick F. Thorne

## Abstract

Loss and overexpression of FAT1 occurs among different cancers with these divergent states equated with tumor suppressor and oncogene activity, respectively. Regarding the latter, FAT1 is highly expressed in a high proportion of human acute leukemias relative to normal blood cells, with evidence pointing to an oncogenic role. We hypothesized that this occurrence represents legacy expression of FAT1 in undefined hematopoietic precursor subsets that is sustained following transformation, predicating a role for FAT1 during normal hematopoiesis. We explored this concept by using the Vavi-Cre strain to construct conditional knockout (cKO) mice where Fat1 expression was deleted at the hemopoietic stem cell stage. Extensive analysis of precursor and mature blood populations using multi-panel flow cytometry revealed no ostensible differences between Fat1 cKO mice and normal littermates. Further functional comparisons involving colony forming unit and competitive bone marrow transplantation assays support the conclusion that Fat1 is dispensable for normal murine haematopoiesis.

## Introduction

Relative to normal hematopoietic cells, FAT1 cadherin is overexpressed in 60% of T-cell and 30% of preB-cell paediatric acute lymphoblastic leukemia (ALL) cases [1]. High FAT1 expression also occurs in half of adult T-ALL cases and in 23% of adult acute myeloid leukemia (AML) cases [2]. Supporting an oncogenic role, FAT1 overexpression is associated with faster relapse in preB-ALL [1], and a unique N-terminal truncated form of FAT1 expressed in T-ALL cooperates with NOTCH1 to accelerate leukemia progression [3].

Alternatively, investigations in different cancers have proposed FAT1 is a tumour suppressor based on its differential expression and mutational status [4]. Indeed, Drosophila *Fat* was originally reported to act as a tumour suppressor [5], and high rates of *FAT1* deletion occur in squamous cell carcinomas (SCC) [6]. In that setting, FAT1-deficient SCC tumours display increased stemness characteristics and experimentally, loss of *Fat1* in a murine model accelerates SCC progression [7].

Here, given the low FAT1 expression in the mature blood cell lineages, we hypothesised that FAT1 expression in leukemia represents legacy expression in undefined haematopoietic precursor subsets that is sustained following transformation. Further, this would predicate a role for FAT1 during haematopoiesis. To address this hypothesis, we generated the first conditional knockout mouse targeting Fat1 in the blood compartment. Extensive phenotypic and functional comparisons employing *in vitro* and *in vivo* approaches indicated no differences between the Fat1 knockout mice and control littermates. These findings provide the definitive conclusion that Fat1 does not contribute to normal hematopoiesis, providing important lessons for deciphering its role during malignant hematopoiesis while also strengthening the concept of FAT1 as a therapeutic target.

## Methods

### Mouse strains and mating strategies

*Fat1*^*fl/fl*^ mice obtained from the laboratory of Professor N. Sibinga contain *Loxp* sites flanking exon 2 of *Fat1. Vav-iCre* transgenic mice (ref) were purchased from Cyagen (#C001019). The *Fat1*^*fl/fl*^ mice were crossed with *Vav-iCre* mice to produce *Vav-iCre;Fat1*^*fl/fl*^ (*Fat1*^*cKO*^). Mice were routinely genotyped using genomic DNA from tail snips with PCR reactions performed with amplicons of 210 and 310 nucleotides, respectively, for the wild-type allele and mutant alleles (forward primer 5′ GTCTGTGTTTGGCCTGAAGACGTA 3′; reverse primer was 5′ AGCACTTCCCAACACCAACATCAA 3′). Both strains were on a C57BL/6-Ly5.2 (Ly5.2, CD45.2) background with both female and male mice aged 6-8 weeks used for experiments together with age and gender matched littermates as controls. Peripheral blood genomic DNA was extracted using a blood/cell/tissue genomic DNA extraction kit (CWBIO), and the recombinantly spliced bands verified by PCR (forward primer 5′ GCTGGCCTGGGTTACGTGAAATT 3′; reverse primer was 5′ CCTAATACACAGCCACC AGCCA 3′). Where indicated, B6.SJL-Ly5.1 (CD45.1) obtained from The Jackson Laboratory were used. All mice were maintained under specific pathogen–free conditions and were used according to the protocols approved by the Institutional Animal Care and Use Committee of Zhengzhou University Medical Center (ZZU-LAC20211210-01).

### Cell and tissue preparation

Tail vein puncture was used for repeated collection of PB with samples immediately introduced into 5 mL FACS tubes containing PBS supplemented with 2% FBS and 5mM EDTA (PBE buffer). The diluted samples were depleted of RBC using Red Blood Cell Lysis Buffer (R1010; Solarbio) according to the manufacturers protocol. BM single cell suspensions were pooled from both femurs, tibias and hips with White Blood Cell Dilution reagent (G3600; Solarbio) used to obtain viable leukocyte counts using a haemocytometer. Whole tissues of interest (i.e., thymus, spleen, and liver) were fixed in formaldehyde for later histological analysis using hematoxylin and eosin staining.

### Complete blood counts

PB was obtained using submandibular vein puncture and immediately introduced into 1.5 mL tubes containing dried EDTA as anticoagulant. Thereafter, samples were analysed using a HEMAVET-950 hematology analyzer (DREW Scientific) with detection parameters set to “mouse”.

### Flow cytometric analyses

PB or BM cell suspensions in PBE buffer were stained on ice for 20 min with the indicated multi-colour antibody panels consisting of fluor-labelled antibodies directed against CD3 (13-0032-82, eBioscience/100206, Biolegend), CD4 (13-0043-82, eBioscience/ 11-0041-82, eBioscience), CD8a (13-0081-82, eBioscience/47-0081-82, eBioscience), CD11b/Mac1 (557657, BD/ 13-0112-82, eBioscience), Gr-1 (108406, Biolegend/ 13-5931-82, eBioscience), B220 (552772, BD/13-0452-82, eBioscience), CD71 (561936, BD), Ter119 (557853, BD/ 13-5921-82, eBioscience), CD45.2 (109828, Biolegend), CD45.1 (110714, Biolegend), CD34 (11-0341-82, eBioscience), CD48 (61-0481-82, eBioscience), Sca-1 (108114, Biolegend), c-Kit/CD117 (560557, BD/47-1171-82, eBioscience/17-1171-82, eBioscience), Flk2/CD135 (46-1351-82, eBioscience/135305, Biolegend), CD16/32 (560540, BD), CD150 (115910, Biolegend), CD127/IL-7Ra (135012, Biolegend), and CD127/IL-7Ra (135012, Biolegend). DAPI (1:2000 of 1 mg/mL stock; Solarbio C0060) was added prior to data acquisition using a FACS AriaIII (BD). Single live cells were analysed with FlowJo software (TreeStar) according to the gating strategies in the corresponding Supplementary materials.

### RNA extraction and quantitative real-time polymerase chain reaction (qRT-PCR)

Total RNA was isolated with an RNA pure Tissue & Cell Kit (CWBIO) or TRIzol™ Reagent (Thermo Fisher) according to the manufacturer’s instructions. cDNA was synthesized with the PrimeScript™ RT Master Mix reverse transcriptase (RR036A; TaKaRa) according to the manufacturer’s instructions with qRT-PCR performed using SYBR Green Master Mix (SparkJade) using a StepOnePlus™ real-time PCR intrument (ThermoFisher). Fat1 levels were measured using forward (5′ GGACCAGCATCGCAAGAGTC 3′) and reverse (5′ GCAAACACGTCAGTTCTGTCTTT 3′) primers with data normalized against actin levels.

### Colony-forming blast unit (CFU) assays

Viable BM cells (2×10^4^) were seeded in 1.1 mL Mouse Methylcellulose Complete Media (M3434; Stem Cell Technologies) in uncoated 6 well plates (3471; Corning) and cultured at 37°C, 5% CO_2_ for 10 days. Colony counts were performed according to standard protocols to identify granulocyte-macrophage (GM), granulocyte-erythroid-macrophage-megakaryocyte (GEMM), and erythroid burst forming unit (BFU-E) colonies. Each sample was performed in triplicate with results reported as average CFU/well.

### Competitive BM transplantation assays

Bone marrow cell isolates obtained from either donor *Vav-iCre;Fat1*^*fl/fl*^ or *Fat1*^*fl/fl*^ mice were combined with isolates from B6.SJL-Ly5.1 (CD45.1) at a 1:1 ratio (5×10^6^ total viable nucleated cells per mL) before intravenous tail vein injection of 1×10^6^ total cells into lethally irradiated (8.5Gy) CD45.2 recipient mice. PB of the recipient mice was then analysed by flow cytometry at the indicated intervals using specific antibodies detecting CD45.1 and CD45.2, respectively. Where indicated, the mice were subjected to irradiation with 5Gy and PB further monitored as indicated.

### Statistical analyses

When parameters followed Gaussian distribution, Student’s t test was used to compare two groups, One-way-ANOVA was used when comparing more than two groups. Data were analyzed using Prism 8.0 (GraphPad Software). In the figures, data are expressed as mean ± standard error (SEM) and significance was set at P < 0.05 (asterisks indicate * P < 0.05, ** P < 0.01, and *** P < 0.001). Sample size ‘n’ indicates biological replicates.

## Results & Discussion

Our primary goal was to address the role of Fat1 in the haematopoietic system using a gene knockout strategy in mice. We first confirmed that the expression profile of Fat1 during murine hematopoiesis was ostensibly like that reported in human subsets (de Bock *et al*, 2012) (Figure S1a-f). Thereafter, given that germline Fat1 deletion is perinatally lethal [8], we generated conditional gene knockout (cKO) mice by crossing Vav-iCre transgenic mice [9] against those bearing a Fat1 conditional (floxed) allele [10] (Fig. 1a). Genotyping identified wildtype, heterozygous and homozygous (cKO) conditionally targeted Fat1 mice (Fig. 1b) with significant reductions in Fat1 mRNA expression shown in bone marrow between wildtype and cKO mice (Fig. 1c). Moreover, the recombinant Fat1 floxed allele was detected in the peripheral blood of cKO mice (Fig. 1d) demonstrating Fat1 loss throughout the hematopoietic system.

**Figure 1.**
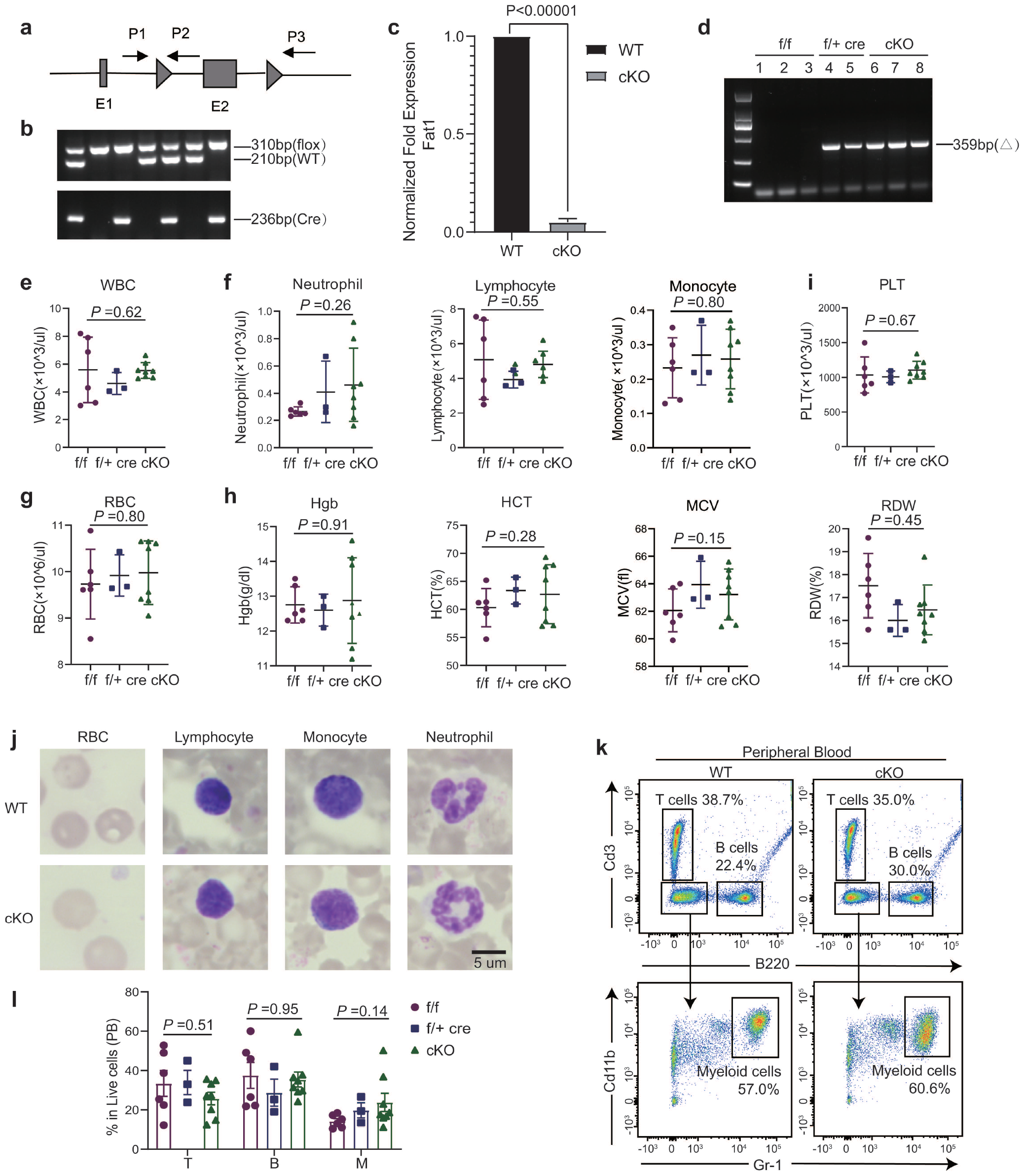
Construction of mice with Vav-iCre-mediated conditional knockout of Fat1 and analysis of mature blood cell populations. (a) Schematic of *Fat1* showing LoxP sites flanking exon 2 (triangles) with arrows denoting genotyping primers (P1-P3). (b) Genotyping of tail gDNA detecting wildtype (P1-P2 product; 210 bp) or floxed alleles (P1-P3 product; 310 bp) along with the presence of Cre recombinase. (c) Comparative qPCR measurement of Fat1 expression in bone marrow isolated from Fat1 f/f (wildtype) and Fat1 f/f Cre (cKO) mice. Bone marrow suspensions were prepared from femurs, tibias and hips before using White Blood Cell Dilution reagent (G3600; Solarbio) to obtain viable leukocyte counts. (d) Genotyping of peripheral blood (PB) from wildtype, f/+ cre (heterozygous) and cKO mice detecting the floxed allele (P1-P3 product; 359 bp). (e-j) PB parameters in Fat1 wildtype, heterozygous and cKO mice reporting counts for white blood cells (WBC) (e), neutrophils, lymphocytes and monocytes (f), total red blood cells (RBC) (g), key RBC parameters, hemoglobin (Hgb), hematocrit (HCT), mean corpuscular volume (MCV), and red blood cell distribution width (RDW) (h) along with blood platelet (PLT) counts (i) and representative cells from blood smears (j). Samples were analysed using a HEMAVET-950 hematology analyzer (DREW Scientific) using “mouse” parameters. (k, l) Representative flow cytometric plots comparing mature T, B and myeloid (M) cells in the PB of Fat1 wildtype and cKO mice (k) along with histogram plots reporting all genotypes (l). PB was collected by tail vein puncture and depleted of RBC using Red Blood Cell Lysis Buffer (R1010; Solarbio). (c, e-h, l) represent findings from 7-week-old mice (n=6 mice/group) except for c (n=3 mice/group), with data reported as mean ± sem. Statistical differences were determined using t-test (two group comparisons) or one way ANOVA (three group comparisons) using GraphPad Prism 8.0 software.

Fat1 cKO mice were born at Mendelian frequencies (Table 1) with homozygous animals displaying normal appearance and growth. No gross phenotypic or histology changes were evident in major hematopoietic organs including the liver, spleen and thymus (Fig. S2a-f), except for a trend of smaller thymus weights in cKO mice (Fig. S2g). Follow-up assessment of intrathymic T-cell development revealed no significant abnormalities (Fig. S2i-k). Moreover, there was no significant differences between the different genotypes for total WBC counts (Fig. 1e), total neutrophil, lymphocyte and monocyte counts (Fig. 1f), RBC counts and other key RBC parameters or platelet counts (Fig. 1g-i). Furthermore, similar cell phenotypes were evident in blood smears (Fig. 1j), while immunotyping assays for T-, B- and myeloid cells showed no changes in mature subsets in the blood or bone marrow associated with Fat1 deficiency (Fig. 1k,l and Fig. S3a-c).

**Table 1.** Expected and observed genotype frequencies of Fat1 f/f (wildtype) and Fat1 f/f Cre (cKO) mice.

| Genotype | *Fat1<sup>fl/fl</sup>* | *Fat1<sup>fl/fl</sup>; iCre* |
| --- | --- | --- |
| Ratio | 1:2 | 1:2 |
| Expected | 33.5 | 33.5 |
| Observed | 34 | 33 |

These findings prompted us to examine if loss of Fat1 affects haematopoietic precursors using LSK-based immunophenotyping strategies (low lineage marker expression (Lin-) together with Sca-1 and c-kit positivity) along with analyses of downstream lineage-committed precursors (Fig. S4a-g). Analyses of bone marrow comparing Fat1 wildtype, heterozygous and cKO mice demonstrated no significant differences between the genotypes comparing LSK, LT-HSC, ST-HSC and MPP populations along with CD150+LT-HSCs [11, 12] (Fig. 2a and Fig. S5a, upper panels). Similarly, the results of a CD150/CD48 immunotyping strategy [13] showed no differences among SLAM-HSCs, SLAM-MPPs, HPC1 and HPC2 populations between Fat1 wildtype and cKO mice (Fig. 2a and Fig. S5a, lower panels). Furthermore, there was no differences for the downstream lineage-committed precursor populations for the myeloid, granulocyte/monocyte, megakaryocyte/erythroid, and lymphoid lineages [14-16] along with proerythroblast/erythroblast subpopulations [17] (Fig. 2c-e and Fig. S5b-d). Colony-forming unit (CFU) assays also found no differences in phenotypes or total colonies derived from Fat1 wildtype and cKO bone marrow progenitors (Fig. 2f, g). Taken together, loss of Fat1 results in no differences in the size or distribution of haematopoietic progenitor pools.

**Figure 2.**
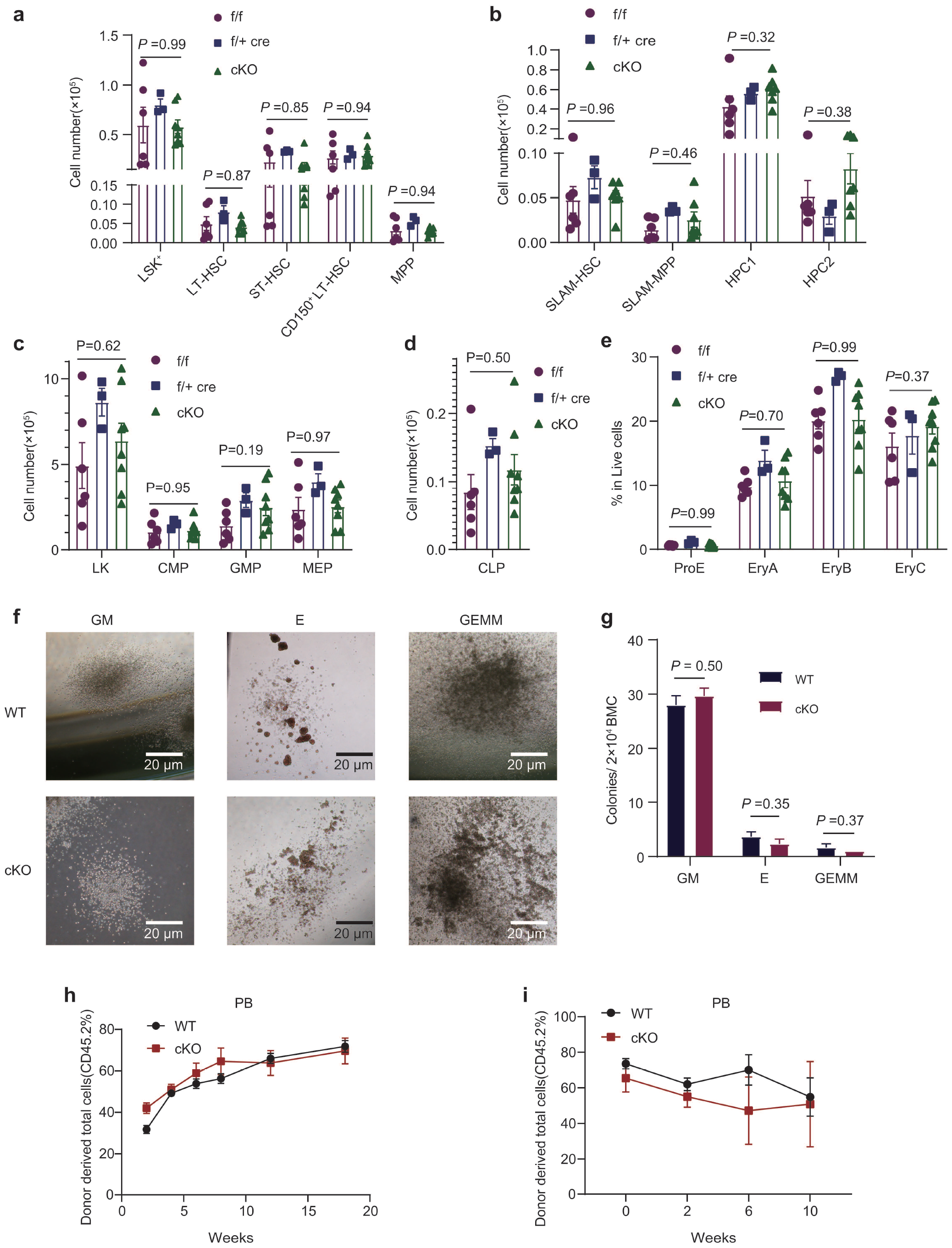
Examination of hematopoietic stem cell maintenance, differentiation, and function after conditional Fat1 knockout. (a) Flow cytometric analyses of hematopoietic precursor subsets in the bone marrow (BM) of Fat1 f/f (wildtype), f/+ Cre (heterozygous) and f/f Cre (cKO) mice using the LSK+ (Lin–Sca1+c-Kit+) definitions: LT-HSC (Lin–Sca1+c-Kit+CD34–Flt3–), ST-HSC (Lin–Sca1+c-Kit+CD34+Flt3–), MPP (Lin–Sca1+c-Kit+CD34+Flt3+) and CD150+ LT-HSC. (b) Flow cytometric analyses of SLAM-defined HSCs as per (a) using the SLAM definitions: HSC (Lin– Sca1+c-Kit+CD150+CD48-), MPP (Lin– Sca1+c-Kit+CD150– CD48–), HPC1 (Lin– Sca1+c-Kit+CD150–CD48+) and HPC2 (Lin– Sca1+c-Kit+CD150+CD48+). (c-e) Flow cytometric analyses of multipotent common myeloid (CMP), megakaryocyte-erythrocyte (MEP) and granulocyte-monocyte progenitors (GMP) (c), common lymphoid progenitors (CLP) (d) and erythropoietic precursors defined using the ProE, EryA, EryB and EryC subsets (e) in the bone marrow of Fat1 wildtype, heterozygous and cKO mice. (f, g) Viable BM cells (2×10^4^) from wildtype and cKO mice were seeded in 1.1 mL Mouse Methylcellulose Complete Media (M3434; Stem Cell Technologies) in uncoated 6 well plates (3471; Corning) and cultured at 37°C, 5% CO_2_ for 10 days. Colony counts were performed according to standard protocols to identify granulocyte-macrophage (GM), granulocyte-erythroid-macrophage-megakaryocyte (GEMM), and erythroid burst forming unit (BFU-E) colonies with results reported as average CFU/well. (h, i) Competitive bone marrow transplantation assays comparing reconstitution potential of CD45.2+ Fat1 replete or cKO donor BM versus CD45.1+ wildtype donor BM. Flow cytometric monitoring of the proportion of CD45.2+ cells in peripheral blood (PB) conducted over 2-18 w (n=7 mice/group; h). Thereafter, mice were subjected to 5Gy irradiation at 20 w and CD45.2+ cells monitored in PB over a further 10 w (n=5 mice/group; i). (a-e) represent findings from 7-week-old male and female mice (n=6 mice/group), (g) (n=3 wells/group). All data reported as mean ± sem with statistical differences determined using t-test (two group comparisons) or one way ANOVA (three group comparisons) using GraphPad Prism 8.0 software.

Next, competitive bone marrow transplantation assays were undertaken to determine if Fat1 loss affected HSC fitness (Fig. S6a). HSCs from the wildtype and Fat1 cKO genotypes displayed near identical reconstitution potential over 2-18 weeks (Fig. 2h), with no major differences in the recovery of myeloid, T-cell or B-cell lineages (Fig. S6b-d). Given the recent link between Fat1 and mitochondrial function [10], we then tested whether Fat1 loss resulted in differences in HSC function under stress conditions using secondary irradiation that elicits DNA damage and free radical generation with subsequent inflammatory responses. However, no differences were observed between wildtype and cKO mice (Fig. 2h and Fig. S6e-h), suggesting Fat1 expression does not contribute to stress recovery.

This study further reinforces that cellular context is crucial for understanding the role of FAT1 in both normal tissues and cancer. In normal development, transgenic Fat1 KO mice uncovered partially penetrant neural defects during embryogenesis while pups developing to term exhibited perinatal lethality [8]. More recently, *Fat1* conditional knockout models uncovered a role for Fat1 in vascular repair via effects on mitochondrial activity [10]. Here, using conditional knockout mice to eliminate Fat1 in HSCs, we show that Fat1 dispensable for haematopoiesis. This in turn infers that the prominent FAT1 expression in acute leukemias is an active oncogenic event, a notion supported by a recent study in T-ALL where many cases show FAT1 promotor hypomethylation [18]. From the broader perspective where Fat1 has often been linked with Hippo pathway regulation in different contexts [19], it is interesting to note that elimination of the Hippo effectors Yap and Taz was similarly dispensable for murine haematopoiesis [20]. An important clinical implication from this study is that ectopic FAT1 expression in leukaemia presents an excellent immunotherapy target, particularly for T-ALL where targeting of common antigens such as CD7 leads to fratricide which could potentially be avoided, although the caveat of FAT1 expression in other tissues needs to be considered.

## Abbreviations

ALL: acute lymphoblastic leukemia
BM: bone marrow
CFU: colony-forming unit
CLP: common lymphoid progenitor
CMP: common myeloid progenitor
(cKO): Conditional knockout
GEMM: granulocyte-erythroid-macrophage-megakaryocyte
GM: granulocyte-macrophage
GMP: granulocyte/monocyte progenitor
HSC: hematopoietic stem cell
MEP: megakaryocyte/erythroid progenitor
MPP: multipotent hematopoietic progenitor
PB: peripheral blood
ProE: proerythroblast
qRT-PCR: quantitative real-time polymerase chain reaction
SLAM: signaling lymphocyte activation molecule

## Authorship

Q.Z. performed most data curation and formal analysis with the assistance from X.Y.H., Y.Y.W., P.P.Z., L.N.C., and R.H.Y. The original draft writing including figures and methods was performed by Q.Z. with R.F.T., C.D.B., Z.M.Z., S.C. and X.D.Z. contribution project conceptualization and oversight, along with reviewing, editing, and revising the final manuscript.

## Acknowledgments

We are indebted to Prof. Nicholas Sibinga and his colleagues (Einstein College of Medicine) for their generous gift of the Fat1 mouse strain. The authors gratefully acknowledge funding from the National Natural Science Foundation of China (81970153). R.F.T., C.D.B., and S.C. conceived the study.

