## Supplementary Figures for "The tumour suppressor Fat1 is dispensable for normal murine hematopoiesis"

### Supplemental figures

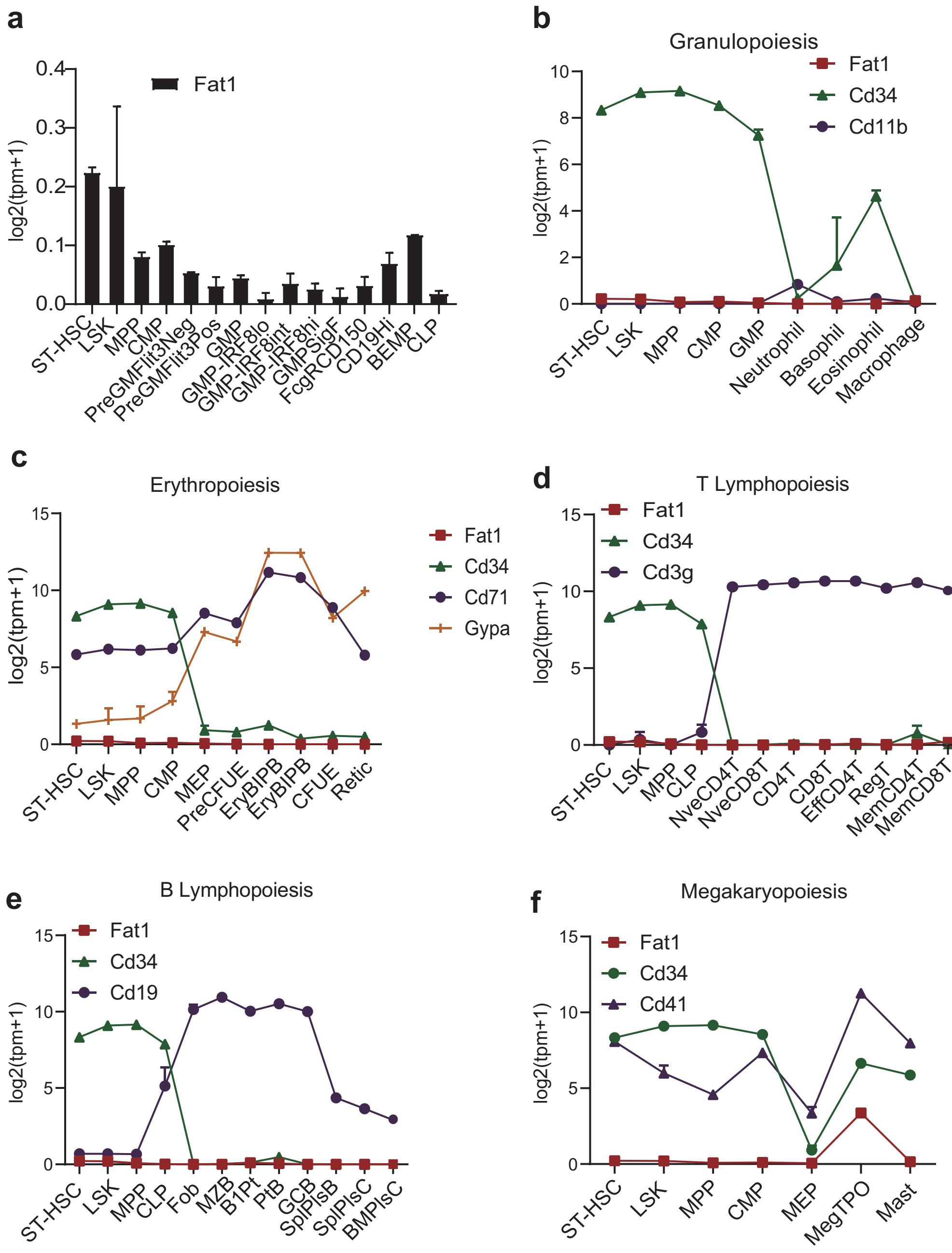
a
Fat1
log2(tpm+1)
0.4
0.3
0.2
0.1
0.0
ST-HSC
LSK
MPP
CMP
PreGMFlit3Neg
PreGMFlit3Pos
GMP
GMP-IRF8lo
GMP-IRF8int
GMP-IRF8hi
GMPSigF
FcgRCD150
CD19Hi
BEMP
CLP
b
Granulopoiesis
Fat1
Cd34
Cd11b
log2(tpm+1)
ST-HSC
LSK
MPP
CMP
GMP
Neutrophil
Basophil
Eosinophil
Macrophage
c
Erythropoiesis
Fat1
Cd34
Cd71
Gypa
log2(tpm+1)
ST-HSC
LSK
MPP
CMP
MEP
PreCFUE
EryBIPB
EryBIPB
CFUE
Retic
d
T Lymphopoiesis
Fat1
Cd34
Cd3g
log2(tpm+1)
ST-HSC
LSK
MPP
CLP
NveCD4T
NveCD8T
CD4T
CD8T
EffCD4T
RegT
MemCD4T
MemCD8T
e
B Lymphopoiesis
Fat1
Cd34
Cd19
log2(tpm+1)
ST-HSC
LSK
MPP
CLP
Fob
MZB
B1Pt
PtB
GCB
SplPlsB
SplPlsC
BMPlsC
f
Megakaryopoiesis
Fat1
Cd34
Cd41
log2(tpm+1)
ST-HSC
LSK
MPP
CMP
MEP
MegTPO
Mast

**Figure S1. Fat1 expression throughout murine hematopoeisis**

(a-f) The relative expression of Fat1 mRNA among defined progenitor cell populations in murine bone marrow (a) along with comparisons with lineage-specific differentiation markers in projections of granulopoiesis (b), erythropoiesis (c), T lymphopoiesis (d), B lymphopoiesis (e) and erythropoiesis (f). Data derived from the Haemosphere database.

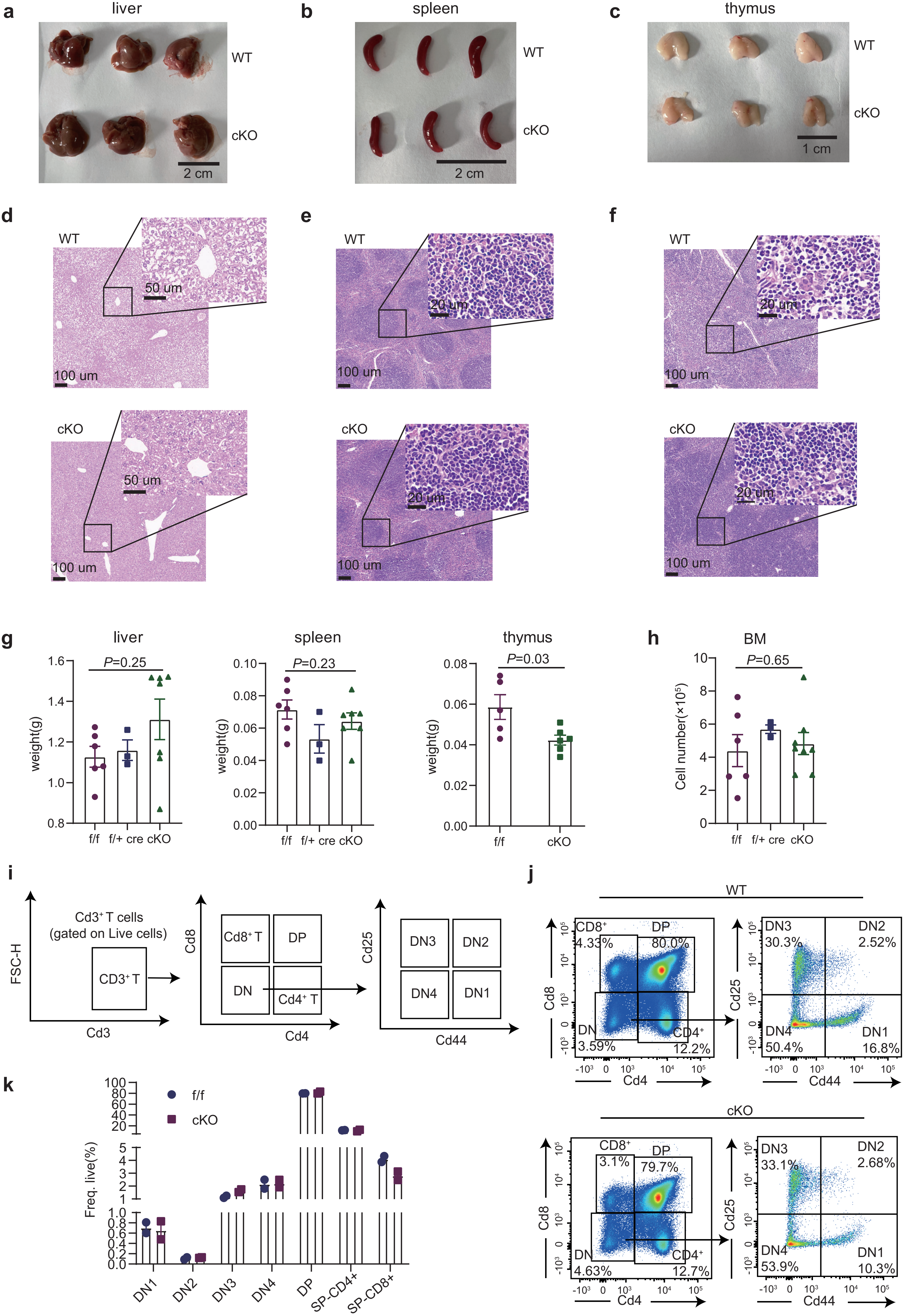

a liver
WT
cKO
2 cm
b spleen
WT
cKO
2 cm
c thymus
WT
cKO
1 cm
d
WT
50 um
100 um
cKO
50 um
100 um
e
WT
20 um
100 um
cKO
20 um
100 um
f
WT
20 um
100 um
cKO
20 um
100 um
g
liver
P=0.25
weight(g)
f/f f/+ cre cKO
spleen
P=0.23
weight(g)
f/f f/+ cre cKO
thymus
P=0.03
weight(g)
f/f cKO
h
BM
P=0.65
Cell number(×10^5)
f/f f/+ cre cKO
i
FSC-H
Cd3+ T cells (gated on Live cells)
CD3+ T
Cd3
Cd8
Cd8+ T
DP
DN
Cd4+ T
Cd4
Cd25
DN3
DN2
DN4
DN1
Cd44
j
WT
CD8+ 4.33%
DP 80.0%
DN 3.59%
CD4+ 12.2%
Cd8
Cd4
DN3 30.3%
DN2 2.52%
DN4 50.4%
DN1 16.8%
Cd25
Cd44
cKO
CD8+ 3.1%
DP 79.7%
DN 4.63%
CD4+ 12.7%
Cd8
Cd4
DN3 33.1%
DN2 2.68%
DN4 53.9%
DN1 10.3%
Cd25
Cd44
k
f/f
cKO
Freq. live(%)
DN1
DN2
DN3
DN4
DP
SP-CD4+
SP-CD8+

**Figure S2. Examination of major haematopoietic organs in Fat1 cKO mice**

(a-f) Gross and histological phenotypes of liver (a, d), spleen (b, e) and thymus (c, f) comparing wildtype (f/f) and cKO (f/f cre) mice.

(g, h) Organ weights (g) and bone marrow (BM) cellularity in the indicated wildtype, heterozygous (f/+ cre) and cKO genotypes. Total BM cells extracted from hips, femurs and tibias.

(i-k) Gating strategy used to define intrathymic steps in T cell development (i) together with representative flow cytometric plots (j) and comparative analyses of wildtype and cKO mice (k).

(g, h, k) represent findings from 7-week-old mice (n=6 mice/group for g, h and n=2 mice/group for k) with data reported as mean ± sem. Statistical differences were determined using t-test (two group comparisons) or one way ANOVA (three group comparisons).

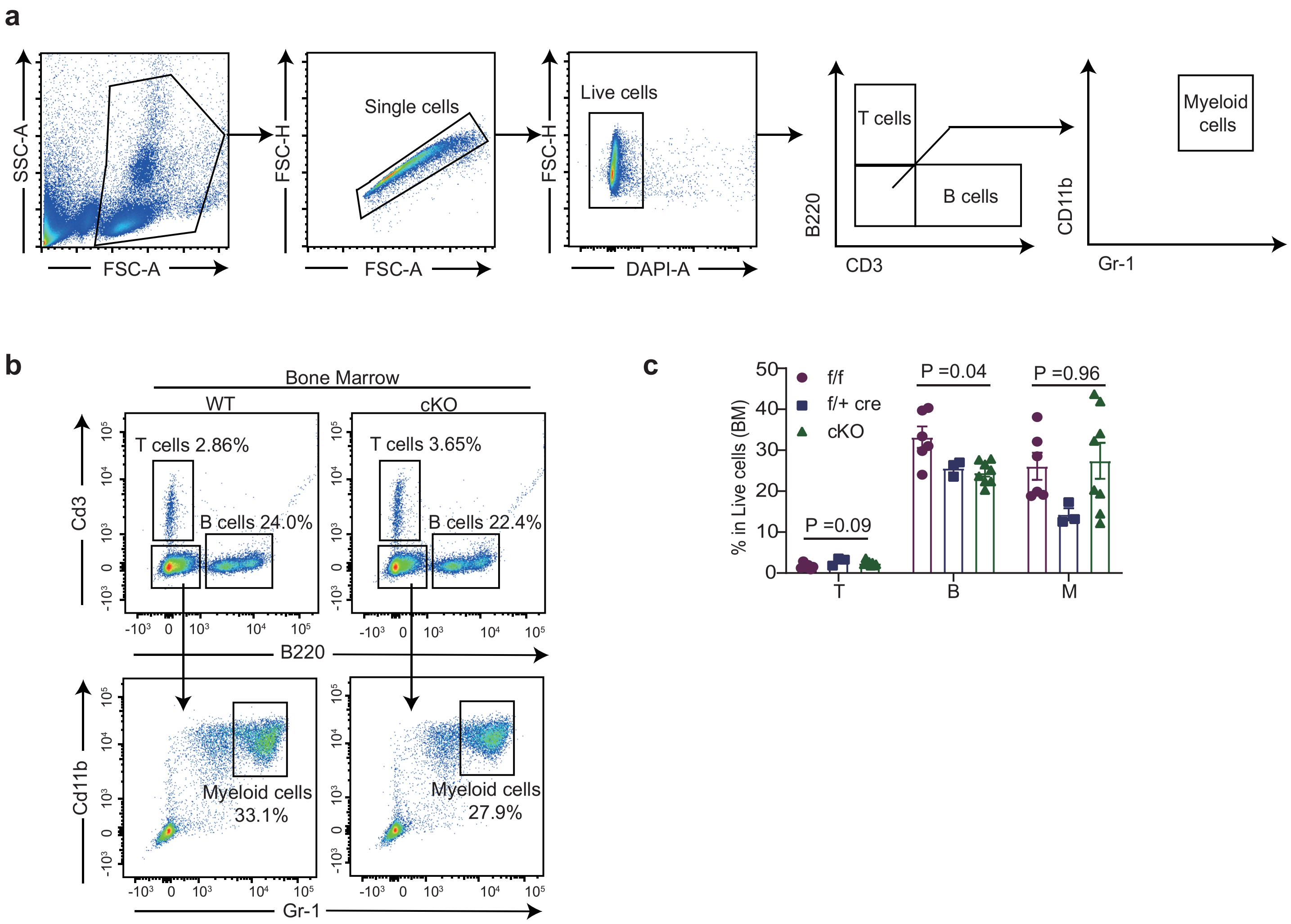
a
SSC-A
FSC-A
FSC-H
Single cells
Live cells
DAPI-A
T cells
B cells
B220
CD3
Myeloid cells
CD11b
Gr-1
b
Bone Marrow
WT
cKO
T cells 2.86%
B cells 24.0%
T cells 3.65%
B cells 22.4%
Cd3
B220
Cd11b
Gr-1
Myeloid cells 33.1%
Myeloid cells 27.9%
c
% in Live cells (BM)
f/f
f/+ cre
cKO
P =0.09
P =0.04
P =0.96
T
B
M

**Figure S3. Analysis of mature hematopoietic cell populations**

(a) Flow cytometric gating strategy outlining the immunophenotyping scheme used to analyse major mature hematopoietic cell lineages (T cell, B cell and myeloid cells) in bone marrow (pictured) or peripheral blood (related to Fig. 1k, l).

(b, c) Representative flow cytometric plots of mature hematopoietic cell lineages in bone marrow comparing wildtype (f/f) and cKO (f/f cre) mice (b) along with cohort analyses of all genotypes including heterozygous (f/+ cre) littermates (c).

(b, c) represent findings from 7-week-old mice (n=6 mice/group) with data reported as mean ± sem. Statistical differences were determined using t-test (two group comparisons) or one way ANOVA (three group comparisons).

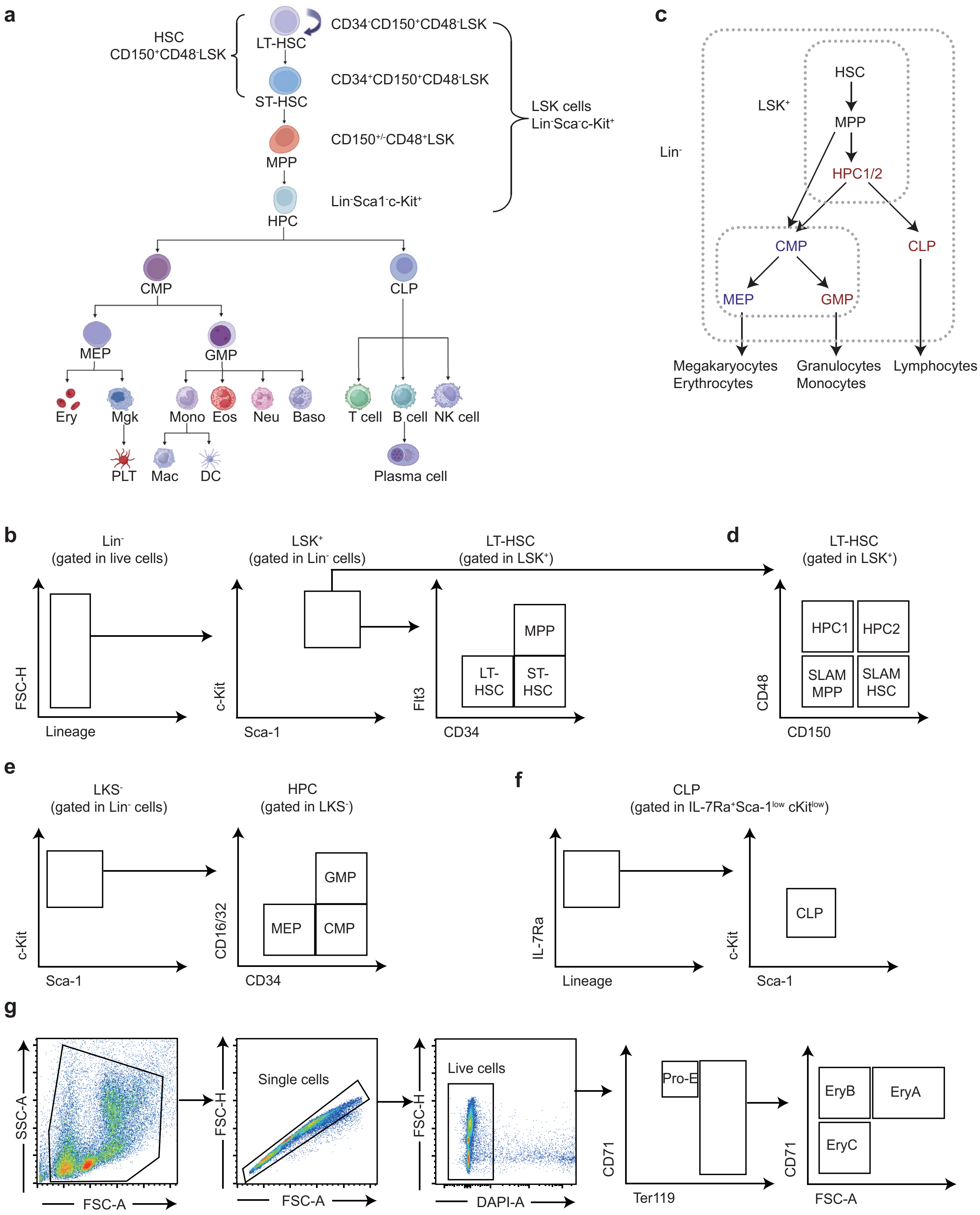
a
HSC
CD150+CD48-LSK
LT-HSC
ST-HSC
MPP
HPC
CD34-CD150+CD48-LSK
CD34+CD150+CD48-LSK
CD150+/-CD48+LSK
Lin-Sca1-c-Kit+
LSK cells
Lin-Sca+c-Kit+
CMP
CLP
MEP
GMP
Ery
Mgk
Mono
Eos
Neu
Baso
T cell
B cell
NK cell
PLT
Mac
DC
Plasma cell
c
HSC
MPP
HPC1/2
LSK+
Lin-
CMP
CLP
MEP
GMP
Megakaryocytes
Erythrocytes
Granulocytes
Monocytes
Lymphocytes
b
Lin-
(gated in live cells)
FSC-H
Lineage
LSK+
(gated in Lin- cells)
c-Kit
Sca-1
LT-HSC
(gated in LSK+)
MPP
LT-HSC
ST-HSC
Flt3
CD34
d
LT-HSC
(gated in LSK+)
HPC1
HPC2
SLAM MPP
SLAM HSC
CD48
CD150
e
LKS-
(gated in Lin- cells)
c-Kit
Sca-1
HPC
(gated in LKS-)
GMP
MEP
CMP
CD16/32
CD34
f
CLP
(gated in IL-7Ra+Sca-1low cKitlow)
IL-7Ra
Lineage
CLP
c-Kit
Sca-1
g
SSC-A
FSC-A
Single cells
FSC-H
FSC-A
Live cells
FSC-H
DAPI-A
Pro-E
CD71
Ter119
EryB
EryA
EryC
CD71
FSC-A

**Figure S4. Analysis definitions of hematopoietic stem cell populations in bone marrow**

(a, b) Illustration depicting hematopoietic stem cell differentiation model based on lineage (Lin) marker expression in concert with Sca-1 and c-kit expression (LSK) (a) with the accompanying flow cytometric gating scheme used to define hierarchical subpopulations of HSCs including LT-HSCs, ST-HSCs, MPPs and HPCs (b) (related to Fig. 2a).

(c, d) Alternative SLAM differentiation model (c) with accompanying flow cytometric gating scheme defining the hierarchical HSC, MPP, HPC1, and HPC2 subpopulations (d) (related to Fig. 2b).

(e, f) Flow cytometric gating scheme defining CMP, MEP, GMP, and CLP subpopulations (e, f) (related to Fig. 2c, d).

(g) Flow cytometric gating scheme used to define erythroid progenitors subsets based on CD71 and Ter119 expression (related to Fig. 2e).

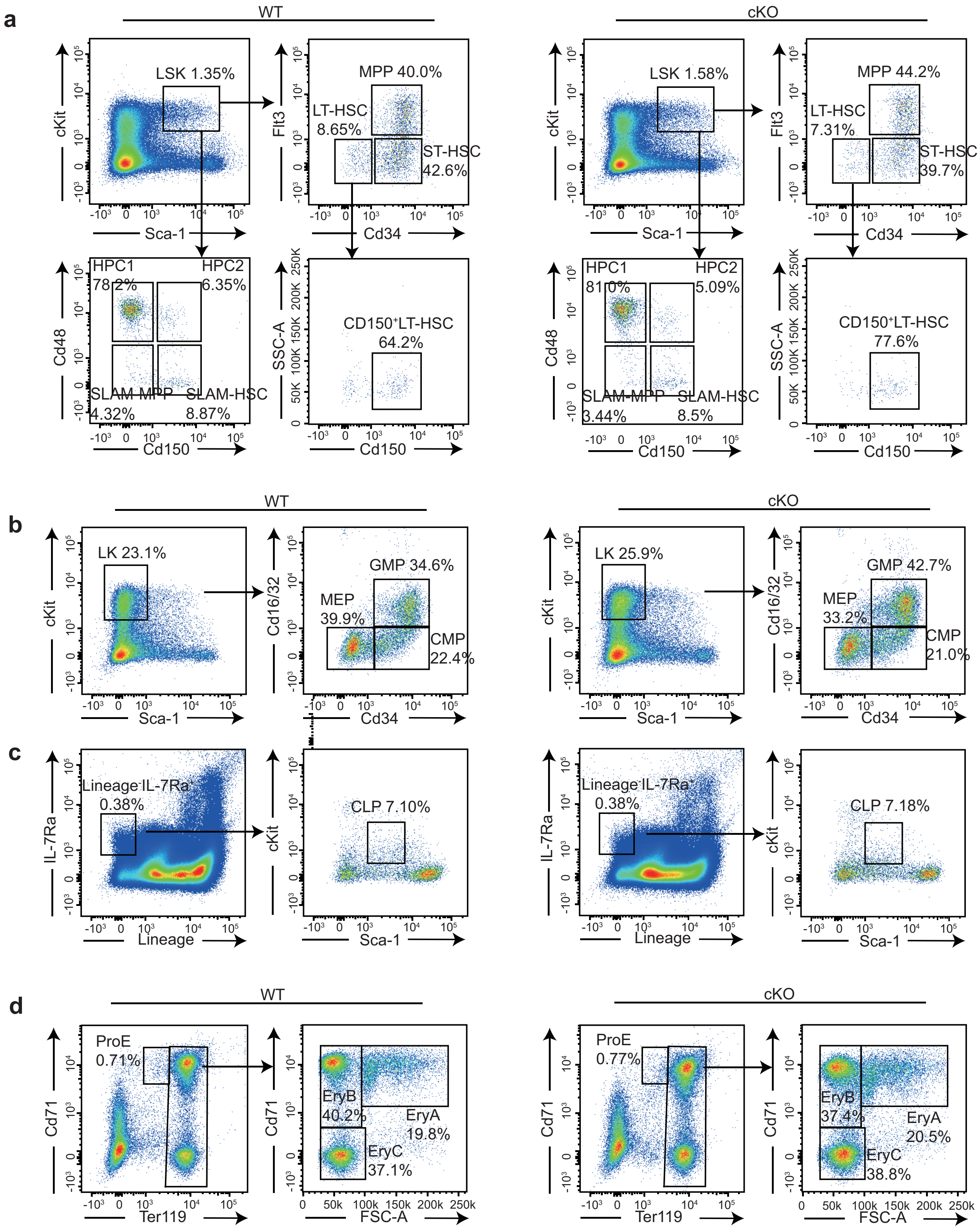
a
WT
cKO
LSK 1.35%
LSK 1.58%
cKit
Sca-1
MPP 40.0%
LT-HSC 8.65%
ST-HSC 42.6%
MPP 44.2%
LT-HSC 7.31%
ST-HSC 39.7%
Flt3
Cd34
HPC1 78.2%
HPC2 6.35%
SLAM-MPP 4.32%
SLAM-HSC 8.87%
HPC1 81.0%
HPC2 5.09%
SLAM-MPP 3.44%
SLAM-HSC 8.5%
Cd48
Cd150
CD150⁺LT-HSC 64.2%
CD150⁺LT-HSC 77.6%
SSC-A
b
LK 23.1%
LK 25.9%
GMP 34.6%
MEP 39.9%
CMP 22.4%
GMP 42.7%
MEP 33.2%
CMP 21.0%
Cd16/32
c
Lineage⁻IL-7Ra⁺ 0.38%
IL-7Ra
Lineage
CLP 7.10%
CLP 7.18%
d
ProE 0.71%
ProE 0.77%
Cd71
Ter119
EryB 40.2%
EryA 19.8%
EryC 37.1%
EryB 37.4%
EryA 20.5%
EryC 38.8%
FSC-A

**Figure S5. Examination of hematopoietic stem cell populations after conditional Fat1 knockout.**

(a) Representative flow cytometric plots of comparing hierarchical hematopoietic precursor populations in the bone marrow of wildtype (f/f) and cKO (f/f cre) mice.

LSK-defined subpopulations represented include multipotent progenitors (MPPs), short-term hematopoietic stem cells (ST-HSCs), long-term hematopoietic stem cells (LT-HSCs) and CD150+ LT-HSCs along with the SLAM-definition subpopulations involving SLAM-HSCs, SLAM-MPPs, and the hematopoietic progenitor cells (HPC) populations 1 and 2 (related to Figs. 2a, b).

(b, c) Representative plots of common myeloid progenitors (CMPs: LSK-CD34highCD16/32high), megakaryocyte/erythroid progenitors (MEPs: LSK-CD34lowCD16/32lowCD150+), granulocyte/monocyte progenitors (GMPs: LSK-CD34middleCD16/32middle) (b) and common lymphoid progenitors (CLPs: IL7Ra+Flt3+ Lin-Sca-1lowc-Kitlow) (c) in the bone marrow of wildtype and cKO mice (related to Figs. 2c, d).

(d) Representative plots of ProE proerythroblast (CD71highTer119intermediate) and the sequential erythroblast subpopulations defined as EryA (CD71highTer119highFSChigh), EryB (CD71highTer119highFSClow) and EryC (CD71lowTer119highFSClow) in the bone marrow of wildtype and cKO mice (related to Fig. 2e).

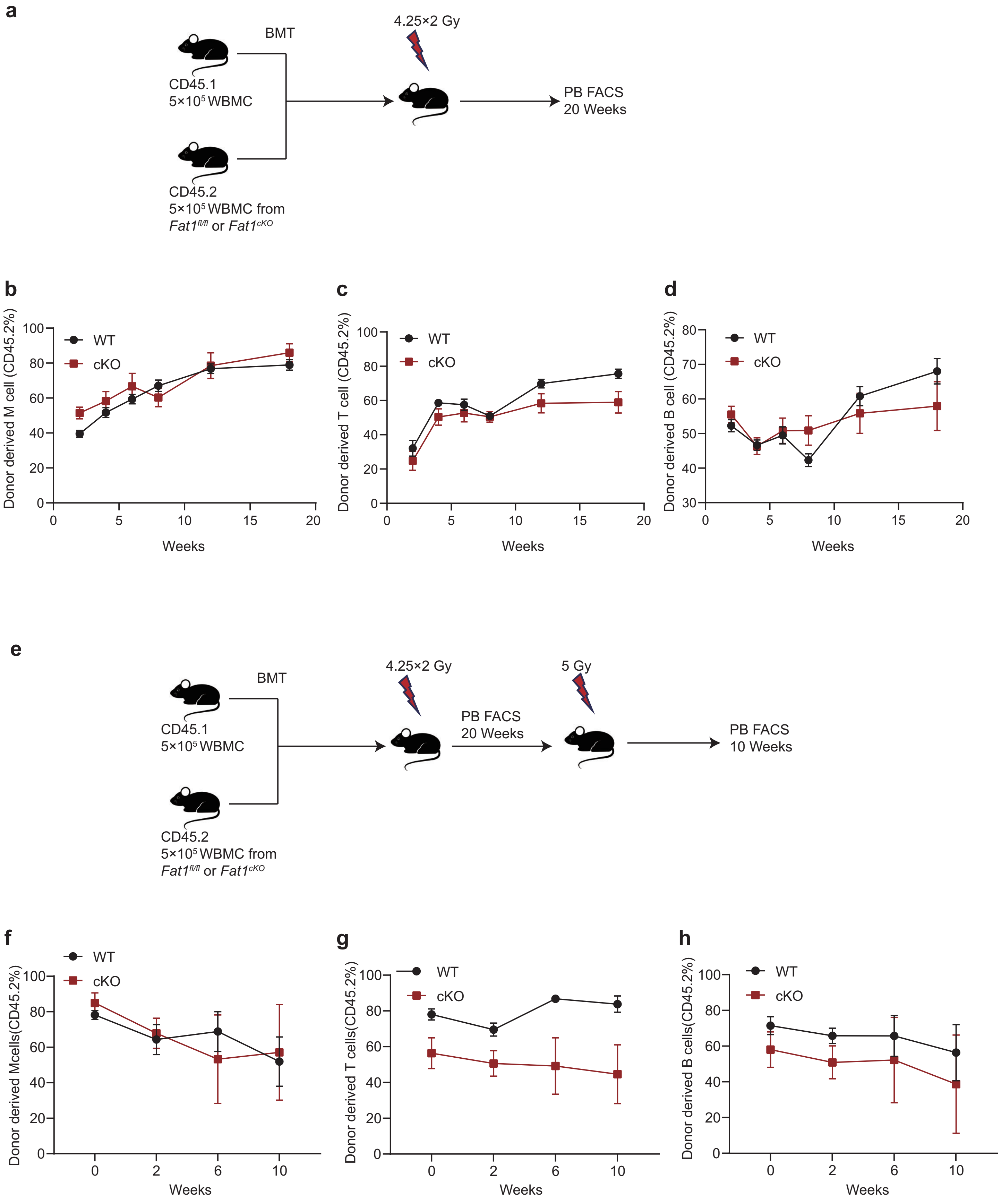

a
BMT
4.25×2 Gy
CD45.1
5×10⁵ WBMC
PB FACS
20 Weeks
CD45.2
5×10⁵ WBMC from
Fat1fl/fl or Fat1cKO
b
c
d
WT
cKO
Donor derived M cell (CD45.2%)
Donor derived T cell (CD45.2%)
Donor derived B cell (CD45.2%)
Weeks
e
BMT
4.25×2 Gy
5 Gy
CD45.1
5×10⁵ WBMC
PB FACS
20 Weeks
PB FACS
10 Weeks
CD45.2
5×10⁵ WBMC from
Fat1fl/fl or Fat1cKO
f
g
h
Donor derived Mcells(CD45.2%)
Donor derived T cells(CD45.2%)
Donor derived B cells(CD45.2%)
Weeks

**Figure S6. In vivo assessment of functional activity of HSCs following conditional Fat1 knockout**

(a, b) Schematic outlining the experimental design of the competitive bone marrow transplantation assay conducted using wildtype C57BL6 (CD45.1) mice in combination with Fat1 wildtype or cKO donors (CD45.2) (a). Flow cytometry determination of the proportion of mature blood myeloid (Gr-1+CD11b+ CD45.2+; E), T (CD3+ CD45.2+; F) and B cell (B220+ CD45.2+; G) populations over 2-18 weeks post-transplantation (b) (n=7/group; related to Fig. 2h).

(c, d) Following 20 weeks recovery, mice from the experiment in (a) were subjected to secondary irradiation (5Gy) and follow-up over a further period of 10 weeks. Experimental scheme (c) and flow cytometric assessment of the proportion of mature blood myeloid, T and B cell populations (d) (n=5/group; related to Fig. 2i).
